# A centrifugation-based method for high-throughput biomaterial separation using magnetic microbeads

**DOI:** 10.1101/2023.04.26.538353

**Authors:** Hiroki Sugishita, Kazunori Hojo, Tetsutaro Hayashi, Itoshi Nikaido, Yukiko Gotoh

## Abstract

Magnetic microbeads are small iron oxide nanoparticles coated with a bioaffinity material that selectively binds to specific biomolecules of interest, enabling their capture and isolation from complex biological samples. Magnetic microbeads are widely used for purification of specific biomolecules in various experiments in molecular biology. However, current methods of manual pipetting to separate supernatants from magnetic microbeads are often inefficient- time-consuming, labor intensive and inaccurate. Furthermore, the use of pipetting robots and liquid handlers specifically designed for multi-well plates can be a cost-prohibitive approach due to the high cost of equipment and disposable supplies. Here, we developed a centrifugation-based method for high-throughput separation of supernatant from magnetic microbeads. To facilitate the centrifugal separation process, we used the 384 transfer plate™ (Watson, Japan) and a magnetic stand equipped with a 384-well magnetic stand, allowing easy handling of several hundred samples and rapid separation of supernatant from magnetic microbeads. The centrifugal force was used to drive the separation of target molecules from the magnetic microbeads, and sample were successfully separated with relatively high recovery rates. Thus, this technology provides a simple, rapid, and cost- and labor-effective biomolecule separation method with potential applications in various fields, including molecular biology, clinical diagnostics, and biotechnology, and is a valuable addition to the existing toolbox of biomolecule separation methods.

## Introduction

Separation (isolation or purification) of a molecule of interest from a mixture of molecules is a fundamental step in biochemical experimental procedures in the life sciences and biotechnology. Separation of target proteins and nucleic acids is carried out, for example, by liquid chromatography, electrophoresis, density gradient centrifugation, ultrafiltration, sedimentation with specific compounds, and various types of column chromatography (ion exchange, hydrophobic, gel filtration, affinity chromatography, etc.). Among them, the use of affinity ligands can be a powerful tool for selective and effective recovery of target molecules. Such affinity ligands include streptavidin (for isolation of biotin-tagged molecules), oligo(dT) (for isolation of mRNA with poly(A) tails), Protein A/G (for isolation of antibodies), and antibody (for isolation of molecules recognized by the antibodies) [1–4]. These affinity ligands are immobilized on solid supports such as sugar- or acrylamide-based polymer resins (produced as hydrated beads of 50-150 µm diameter) and magnetic microbeads (consisting of iron oxide nanoparticles) [4–7]. Polymer resins are generally not suitable for batch separation of target molecules from particulate materials which are abundantly present in crude samples such as cell extracts, given that most batch separation methods require gravity-based sedimentation which precipitates both resins and particulate materials. By contrast, magnetic microbeads can be separated from particulate materials by controlling the angle of the magnetic field away from that of the gravity field. Separating magnetic beads from gravity-based precipitates also has the advantage of effectively removing the washing solutions, by reducing the residual solution after removal. Because of these advantages, the magnetic separation procedure can be simple and easy (given that preclearing of particulate materials is unnecessary) as well as efficient. In addition, magnetic separation procedures are relatively gentle to target molecules compared to conventional column chromatography techniques because of lower shear forces and higher protein concentration during the separation process [7,8]. Furthermore, collection of target molecules by affinity ligands such as by magnetic beads is suitable for yielding high concentration of target molecules even in a small volume, unlike conventional column chromatography which often leads to a large volume of diluted target molecules.

Thus, magnetic microbeads have become a commonly used tool for the isolation of target molecules using affinity ligands [4,7,9–14]. However, some problems remained when using magnetic microbeads in existing manual protocols, especially in experiments with small volume and large numbers of samples. First, in principle, the tension of the liquid solution at the surface of the magnetic beads hampers the removal of the unbound solution by the earth’s gravity [15]. This can be especially problematic when the volume of the solution is small relative to the volume of the magnetic beads, which can lead to inefficient separation of target molecules from unbound molecules as well as to large variation in recovery depending on the amount of solution remaining on the surface of magnetic beads. Second, manual pipetting to remove unbound solutions is labor intensive and time consuming, especially when the number of samples is large, and inaccurate (variable) and inefficient, especially when the volume of samples is small. Third, manual pipetting requires pipette tips, tubes and other disposable parts, which can be costly and generate a significant amount of waste in large scale experiments. These issues of manual pipetting can hinder high-throughput analyses using magnetic beads. The use of liquid handlers designed for 384-well plates can speed up the process (for example, Viaflo 96/384 (INTEGRA Biosciences), epMotion® 96 (Eppendorf), apricot S3 (SPT Labtech)), but these problems remain and a better method for separation of magnetic beads would be desirable.

In this study, we describe a centrifugation-based method for separation of magnetic beads from unbound solutions with the use of the 384 transfer plate™ (WATSON Bio Lab). This method is expected to solve at least some of the existing problems described above, and will enable us to isolate molecules of interest from complex biological samples in a simple/easy and cost-and labor-effective way.

## Results

### Investigation of conditions for efficient recovery of magnetic beads by the centrifugation-based method

We aimed to develop a simple method for high-throughput separation of magnetic beads from supernatant fluid (liquid) using a 384-well plate. For a plate centrifugation, centrifugation is usually performed to collect materials at the bottom of the plate. However, being inspired by the protocol for Quartz-Seq2 [16] which includes a centrifugation step with the plate upside down to collect the materials (liquid solutions) in all wells at once, we hypothesized that centrifugation of the plate with inverted (upside-down) orientation while keeping magnetic beads at the bottom of the plate by a magnetic stand may separate supernatant fluid from the beads. In order to collect the supernatant fluid in each well of the 384-well plate, we used the 384 transfer plate™ (WATSON Bio Lab), which consists of 384 funnels corresponding to the 384-well plate, for transferring the liquid in each well of a 384-well plate to another 384-well plate through a funnel by centrifugation (Fig. 1A-D). To test this idea, we first investigated the conditions under which magnetic beads are effectively trapped on the magnetic stand during centrifugation. We used different type of micromagnetic beads (Dynabeads™ MyOne Streptavidin T1, Dynabeads™ M-280 Streptavidin and Sera-Mag Select™) with a magnetic stand designed for 384-well plate (BiT-Mag384™, Sanplatec). The plate centrifugation was performed with PlateSpin3™ (KUBOTA) using the rapid mode for 30 sec. We first used Dynabeads™ MyOne Streptavidin T1 (Thermo fisher, 1.0 μm diameter) as well as Dynabeads™ M-280 Streptavidin (Thermo fisher, 2.8 µm diameter, 2 µl beads per well), and examined the recovery rate of the beads after centrifugation at different centrifugal force. The recovery rate of the beads at the bottom of the plate was estimated by the amount of beads detected in the pooled supernatant fluids from 16 wells after centrifugation with the turbidity at 630 nm measured by a spectrophotometer. As expected, the unrecovered rate of magnetic beads found in the supernatant fluid became greater by increasing the centrifugal force especially to 300 x *g* (Fig.1E, F). We then used the Sera-Mag Select™ micromagnetic beads (Cytiva, 1.0 µm diameter, 5 µl beads per well), which are commonly used for DNA purification. We again observed that the unrecovered rate of the beads became greater by increasing the centrifugal force (Fig. 1G). Compared to manual pipetting, the centrifugation-based method showed lower recovery rates with the use of Dynabeads™ MyOne Streptavidin T1 and Sera-Mag™ Select micromagnetic beads and similar recovery rates with the use of Dynabeads™□ M-280 Streptavidin at 50-200 x *g* under the conditions indicated (Fig. 1E-G).

**Figure 1.**
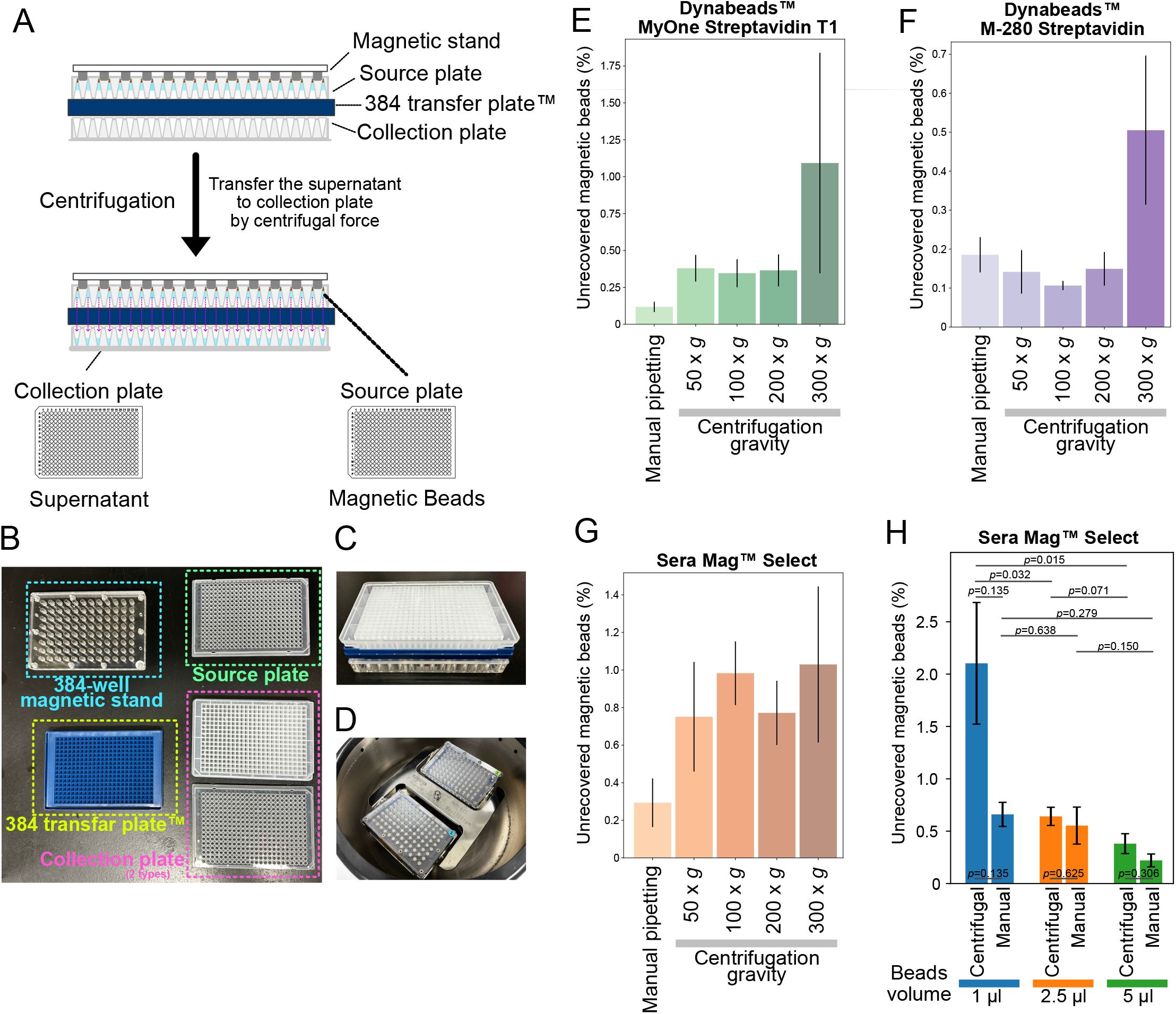
Conditions for recovering magnetic beads using the centrifugation-based method. **A**. Schematic illustration of the centrifugation-based method for high-throughput biomaterial separation using magnetic microbeads. **B**. Pictures of instruments for this centrifugation-based method. **C**. Image of magnetic stand combined with source plate, 384 transfer plate™ and collection plate. **D**. Picture of the setup in the centrifuge machine for the centrifugation-based method. **E-G**. Bar graphs showing the recovery rate in the transferred supernatant of each magnetic microbead (Dynabeads™ MyOne™ Streptavidin T1, Dynabeads™ M-280 Streptavidin, and Sera-Mag™ Select) at different centrifugal force conditions or by manual pipetting. The recovery rate was calculated from the turbidity at 630 nm measured with a spectrophotometer. The turbidity of the supernatant is shown as a ratio to that of the bead-containing solution before centrifugation. Error bars represent SEM (n = 4). **H**. Bar graphs showing the recovery rate in transferred supernatant of Sera-Mag™ Select in different volumes of bead condition by centrifugation-based method or manual pipetting. Error bars represent SEM (n = 4). *p*-value indicates significant differences calculated by Student’s *t*-test (one-tailed).

We then examined whether the amount of beads affects the recovery rate of the beads. We indeed found that the unrecovered rate of Sera-Mag™ Select beads after centrifugation at 200 x *g* was significantly increased when the volume of the beads was reduced from 5 µl to 2.5 µl and 1 µl, which was also the case for manual pipetting (Fig 1H). This indicates that the volume of the Sera-Mag™ Select beads should be greater than 5 µl under the condition used.

### Investigation of conditions for efficient recovery of supernatants by the centrifugation-based method

Next, we estimated the recovery rate of the supernatant fluids after centrifugation at different centrifugal force by measuring the amount of liquid solution transferred to the collection plate (Fig. 2A). In order to measure small amounts of liquid solution, we used DNA-containing solution as a starting material, diluted DNA in the collection plate with 50 µl of TE solution per well and measured its concentration by Qubit™ fluorometer (Fig. 2A). As expected, with the use of Dynabeads™ MyOne Streptavidin T1, the concentration of DNA (indicative of the volume of liquid solution transferred to the collection plate) became slightly greater by increasing the centrifugal force (Fig. 2A). In particular, centrifugation at 200 or 300 x *g* for 30 sec yielded relatively high recovery rates (over 90%) of the supernatant fluid, which were comparable to the separation methods with manual pipetting or the usage of a 96-well liquid handler (Fig. 2A). Given that the recovery rate of the magnetic beads was reduced by centrifugal force greater than 300 x *g* (Fig. 1E), centrifugation at 200 x *g* for 30 sec would be optimal for the centrifugal separation of Dynabeads™ Streptavidin T1 and supernatant fluids among the conditions tested. A similar tendency was found for the use of Dynabeads™□ M-280 Streptavidin and Sera-Mag™ Select beads, but the recovery rate of supernatant fluids tended to be slightly higher and lower than Dynabeads™ Streptavidin T1, respectively, in the range of 50 to 200 x *g* (Fig. 2B, 2C). Given that different magnetic beads behave differently in the recovery rate of the beads and the supernatants, centrifugal conditions should be optimized for each type of the magnetic beads.

**Figure 2.**
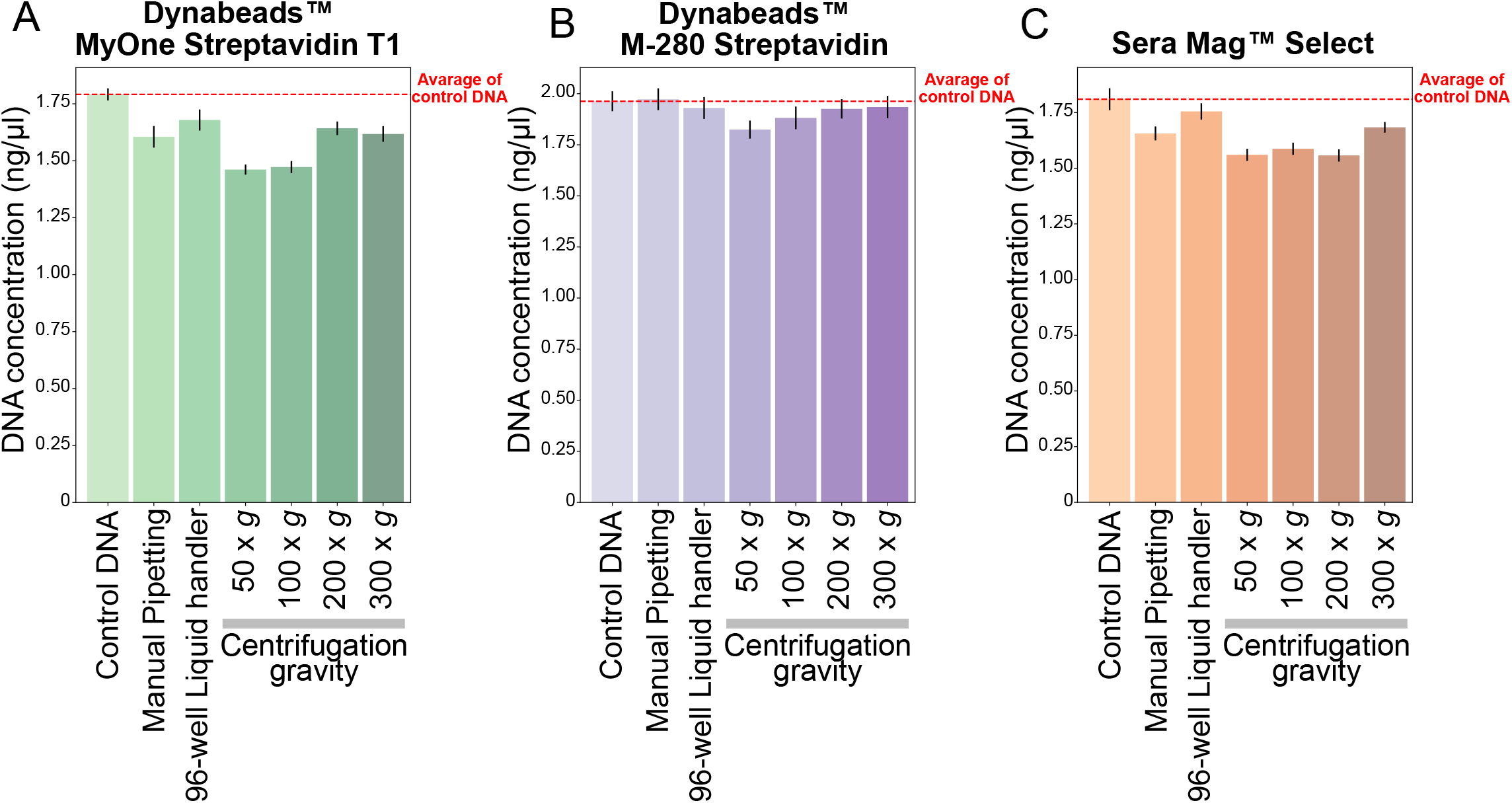
Conditions for recovering supernatants using the centrifugation-based method. **A-C**. Bar graphs showing the DNA concentration (ng/µl) in transferred supernatant DNA diluted with 50 µl of TE solution to measure the volume of liquid collected by the centrifugation-based method at different centrifugal force conditions or by manual pipetting. Each magnetic microbead (Dynabeads™ MyOne™ Streptavidin T1, Dynabeads™ M-280 Streptavidin and Sera-Mag™ Select) was resuspended in DNA after washing, and the DNA solution recovered by centrifugation using a magnetic stand and 384 transfer plate™ was diluted with 50 µl of TE solution and measured the DNA concentration by Qubit™ Flex Fluorometer (Thermo). Error bars represent SEM (n = 8).

### Recovery of DNA in a 384-well plate using the centrifugation-based method with magnetic beads

Finally, to demonstrate the practical utility of this method, we performed DNA purification using Sera-Mag™ Select magnetic beads and a 384-well plate. The DNA purification process involves the following steps: mixing DNA and beads, capturing the beads using a magnet, removing the supernatant (unbound fraction), washing with 85% ethanol three times, drying the beads, eluting the DNA, and collecting the eluate. The centrifugation-based method can be applied not only for the recovery of the eluate but also for the removal of the supernatant, washing with 85% ethanol, and drying of the beads. We thus purified the lambda DNA (Qubit™ 1X dsDNA HS Assay Lambda Standard) for demonstration. We applied a solution of lambda DNA (5 ng/µl) to the wells of the plate only at the positions indicated in Fig. 3A. Subsequently, we added 1.8x volume of Sera-Mag™ Select (3.6 µl) and mixed it by plate mixer (ThermoMixer® C, Eppendorf) after sealing the plate with a lid. We then performed the procedure as shown in Fig. 3A and finally eluted the DNA with 5 µl of TE solution. The eluted solution was collected using a 384 transfer plate™. Lambda DNA was successfully recovered in all wells where it was applied (Fig. 3B). Both manual pipetting method and the centrifugation-based method yielded an average recovery rate of about 50-60%, with the centrifugation-based method showing a slightly higher yield (Fig. 3C). Furthermore, no contaminated DNA was detected in the wells to which TE solution alone was added. This result demonstrated that the centrifugation-based methos with magnetic beads is applicable to high-throughput applications, such as for DNA purification, using a 384-well plate.

**Figure 3.**
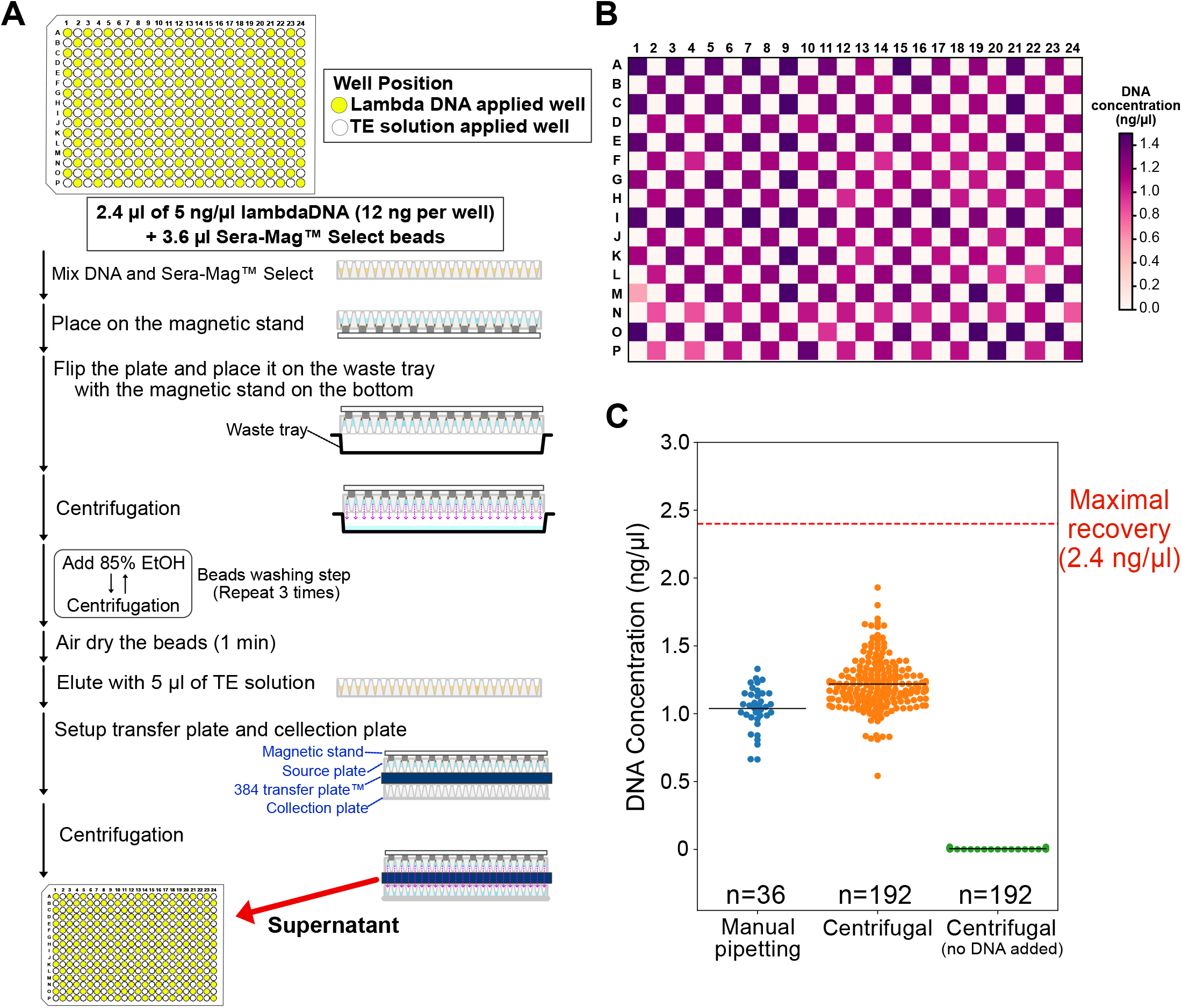
Recovery of DNA in 384-well plates using the centrifugation-based method with magnetic beads. **A**. Schematic illustration of DNA purification using Sera-Mag™ Select beads by centrifugation-based method. **B**. Heatmap showing the DNA concentration of the supernatant after DNA purification in a 384-well PCR plate. The position of the applied DNA was shown in Fig. 3A. **C**. Swarmplot showing the DNA concentration of each well in the centrifugation-based method and manual pipetting experiment.

## Discussion

In this study, we developed a new method for separation of micromagnetic beads and supernatant solution by centrifugation with the use of the 384 transfer plate™ (WATSON Bio Lab) and the 384-well magnetic stand. This method is simpler (easier), faster and cost-effective, which enables simultaneous processing of multiple samples without the need for complex and expensive liquid handling machines. We demonstrated that this centrifugation-based method is applicable to different types of magnetic bead types, namely, Sera-Mag Select™□, Dynabeads™ MyOne Streptavidin T1, and Dynabeads™ M-280 Streptavidin. Given these advantages, this centrifugation-based method appears to be suitable for high-throughput operation of biomolecule separation in a wide range of practical experimental settings.

In this study, we obtained comparable recovery rates of magnetic beads and supernatant fluids with the newly-developed centrifugation-based method and with the conventional manual pipetting method. This is due to the relatively weak magnetic force of the magnetic beads and the magnetic stand we used in this study. Namely, although efficient recovery of the magnetic beads and the supernatant fluid by centrifugation requires strong magnetic force sufficient to withstand the centrifugal force generated by high-speed centrifugation. Therefore, the use of a stronger magnetic stand would increase the recovery rate and may make this separation method more efficient and reliable than conventional manual pipetting method.

In particular, <u>the use of our method for DNA purification is highly applicable in NGS applications, as it allows for the scaling up of samples into the 384-well plate format. This method can also simplify the DNA purification steps required for plate-based single cell RNA-sequencing protocols such as RamDA-Seq, Smart-Seq2/3, and FLASH-Seq [17–21], which typically recommend liquid handling machines designed for 384-well plates</u>. Our method offers a more convenient and efficient solution, with higher recovery rates and <u>the potential for improved final concentrations and yields by reducing the amount of elution volume</u>.

One of the key parts of this method is the usage of the 384 transfer plate™ (WATSON Bio Lab). This plate was designed for the transfer of solutions from one 384-well plate to another by centrifugation. We demonstrated that this plate can be effectively used in conjunction with magnetic beads and a magnetic stand to separate biomaterials. However, it is important to note that the 384 transfer plate™ cannot be for liquid solutions containing oil or high concentrations of surfactants, which results in leakage between the wells. The plate is made of polypropylene, but development of a plate with a low adsorption of biomaterials regardless of liquid properties would improve the versatility and yield of this method.

Overall, our method offers a simpler and more cost-effective approach to high-throughput magnetic bead purification using a 384-well plate, which has the potential to facilitate large-scale screening and analysis in a variety of fields.

## Methods

### Centrifugal separation of magnetic microbeads from supernatant fluids

The magnetic bead (i.e. Dynabeads™ MyOne™ Streptavidin T1 (65601, Thermo), Dynabeads™ M-280 Streptavidin (11205D, Thermo), Sera-Mag™ Select (29343052, Cytiva)) suspension solution was dispensed evenly into the wells of a 384-well PCR plate (PCR384C, Corning) and placed on a magnetic stand for 384-well plate (BiT-Mag384™, Sanplatec) for more than 5 min. Then, a 384 transfer plate™ (1859-384S, WATSON Bio Lab) and a collection plate, which can be either a 384-well PCR plate (PCR384C, Corning) or an Edge-less 384-well Plate™ (1825-384E-110, WATSON Bio Lab), were stacked so that each funnel-shaped pore covered each well. The stacked plates were taped with the collection plate at the bottom, and placed in a plate centrifuge (PlateSpin3™, KUBOTA) with a dummy of the same configuration set as a balance. Centrifugation was performed in rapid mode at different conditions (50 x *g*, 100 x *g*, 200 x *g*, 300 x *g*) for 30 sec.

### Measurement of magnetic bead recovery rate after centrifugation

To measure the recovery rate of magnetic beads in the centrifugation-based method, 2 µl/well of Dynabeads™ MyOne™ Streptavidin T1 (65601, Thermo) or Dynabeads™ M-280 Streptavidin (11205D, Thermo) beads were washed twice with bead wash solution (10 mM Tris-HCl [pH 7. 5], 1 mM EDTA, 2 M NaCl, 0.1% Triton X-100) and then resuspended in 10 µl/well TE solution (10 mM Tris-HCl [pH 8.0], 1 mM EDTA). For Sera-Mag™ Select beads (29343052, Cytiva), 5 µl of beads were added to a 1.5 ml tube and the beads were washed twice with TE solution and then resuspended in 10 µl of TE solution. Next, 10 µl of the bead suspension was added to each of 16 wells in a 384-well plate, which were then centrifuged at 50 x g, 100 x g, 200 x g, or 300 x g for 30 sec. The supernatant was transferred to a collection plate and the turbidity was measured at a wavelength of 630 nm using a plate reader (ARVO X5, PerkinElmer) with a setting of 0.1 sec. For manual pipetting, 10 µl of bead suspension was added to each of 16 wells in a 384-well plate, and the supernatant was carefully collected using a 16-channel manual pipette (INTEGRA Biosciences, 3042). The amount of magnetic beads found in the supernatant was quantified by comparing the measurements to a standard curve.

### Measuring the recovery rate of supernatants after centrifugation

Due to the difficulty of directly measuring small volumes of liquid in the 384-well plate format, we mixed DNA solution into the magnetic beads and separated the supernatant into a collection plate by centrifugation. We then used a large volume of liquid to dilute the sample and estimated the recovery rate by measuring its DNA concentration. To estimate the recovery rate of the supernatant, 2 µl/well of Dynabeads™ MyOne™ Streptavidin T1 (65601, Thermo) or Dynabeads™ M-280 Streptavidin (11205D, Thermo) beads were washed twice with bead wash solution (10 mM Tris-HCl pH 7. 5, 1 mM EDTA, 2 M NaCl, 0.1%Triton X) and then resuspended in lambda DNA solution (diluted to approximately 20 ng/µL in TE solution, 3010, Takara). For the Sera-Mag™ Select (29343052, Cytiva) bead assay, 5 µl/well of beads were washed twice with TE solution and then resuspended in lambda DNA solution (diluted to approximately 20 ng/µL in TE solution, 3010, Takara). Then, 5 µl of each bead suspension was added to 8 wells of a 384-well plate and centrifuged at 50 x g, 100 x g, 200 x g, or 300 x g for 30 sec. The supernatant was transferred to an Edge-less 384-well plate™ (1825-384E-110, WATSON Bio Lab) as a collection plate and DNA recovery was estimated by measuring its concentration using 1x Qubit™ 1X dsDNA HS Assay Kit (Q33231, Thermo) and Qubit™ Flex Fluorometer (Thermo) after dilution with 50 µl TE solution. As a control, 5 µl of DNA supernatant after bead suspension was diluted with 50 µl TE solution. To compare results with the centrifugation-based method, DNA was also collected by manual pipetting and 96-well liquid handling. For manual pipetting, the supernatant was carefully collected using a 16-channel pipette (INTEGRA Biosciences, 3042). For 96-well liquid handling, supernatant was gently collected using a 96-channel pipette head (INTEGRA Biosciences, 6101) on VIAFLO 384™ (INTEGRA Biosciences).

### DNA purification using a 384-well plate and magnetic beads with the centrifugation-based method

For high-throughput DNA purification using magnetic beads and a 384-well plate, we used the centrifugation-based method with Sera-Mag Select. Lambda DNA (Qubit™ 1X dsDNA HS Assay Lambda Standard) was diluted to 5 ng/µl in TE solution. We added 2.4 µl of the diluted lambda DNA (5 ng/µl) and 3.6 µl of Sera-Mag™ Select (1.8x volume) to half of the wells (192 wells, see position in Fig. 3A) of the 384-well PCR plate (PCR384C, Corning). To confirm the absence of well-to-well contamination, 2.4 µl TE solution and 3.6 µl Sera-Mag™ Select (1.8x volume) were added to the other half of the wells (192 wells, see position in Fig. 3A). The plate was sealed with a lid, mixed it by plate mixer (ThermoMixer® C, Eppendorf) at 2000 rpm for 1 min at room temperature and incubated for 5 min. After incubation, the plate was placed on a magnetic stand for 5 min to capture the beads. The plate was then inverted and centrifuged at 100 x g for 30 sec to remove the supernatant (unbound fraction) into the waste tray. To wash the beads, 85% ethanol was added to each well and the plate was centrifuged at 100 x g for 30 sec. This washing step was repeated three times and the beads were dried for 1 min after the last wash. Elution was performed by adding 5 µl of TE solution to each well, and the plate was vortex mixed and incubated at room temperature for 5 min. The eluate was collected in a collection plate using a 384 384 transfer plate™ (1859-384S, WATSON Bio Lab), and the DNA concentration was measured using Qubit™ 1X dsDNA HS Assay. For manual pipetting, 2.4 µl of diluted lambda DNA (5 ng/µl) and 3.6 µl of Sera-Mag™ Select (1.8x volume) were added to 36 wells of a 384-well PCR plate (PCR384C, Corning). The plate was sealed with a lid, mixed it by plate mixer (ThermoMixer® C, Eppendorf) at 2000 rpm for 1 min at room temperature and incubated for 5 min. After incubation, the plate was placed on a magnetic stand for 5 min to capture the beads. The supernatant was removed from each well using a 16-channel pipette (INTEGRA Biosciences, 3042) and 85% ethanol was added to wash the beads. The washing step was repeated three times, and the beads were dried for 1 min after the final wash. Elution was performed by adding 5 µl of TE solution to each well, and the plate was vortex mixed and incubated at room temperature for 5 min. The eluate was transferred to a new 384-well PCR plate (PCR384C, Corning) using a 16-channel pipette (INTEGRA Biosciences, 3042), and the DNA concentration was measured using Qubit™ 1X dsDNA HS Assay.

## Declaration of interests

I.N. and T.H. have been involved in the development of the 384 transfer plate™ (Watson) and BiT-Mag384™ (Sanplatec) and had a previous technology consulting agreement for the 384 transfer plate™ (Watson) and a current consulting agreement for BiT-Mag384™ (Sanplatec).

## Author contribution

Conception: H.S, Y.G, Investigation: H.S. and K.H, Writing: H.S. and Y.G, Funding acquisition: H.S, Y.G., I.N., Resources: T.H and I.N. Supervision: H.S. and Y.G.

## Acknowledgments

This research was supported by AMED under grant number JP23wm0525035 to H.S. ; by AMED-CREST under grant number (JP23gm1310004) to Y.G. ; by KAKENHI grants from the Ministry of Education, Culture, Sports, Science, and Technology of Japan and the Japan Society for the Promotion of Science (JP22K15118 to H.S., JP22H00431 to Y.G.); by the Uehara Memorial Foundation, and by the International Research Center for Neurointelligence (WPI-IRCN), The University of Tokyo Institutes for Advanced Study to H.S and Y.G.. This research was partially supported by JST CREST (JPMJCR16G3, JPMJCR1926 and JPMJCR21N6), AMED (JP21bm0404073, JP22fk0210093, and JP21fk0210093), and the Medical Research Center Initiative for High Depth Omics and Nanken-Kyoten, TMDU to I.N.

## REFERENCES

[1] B.V. Ayyar, S. Arora, C. Murphy, R. O’Kennedy, Affinity chromatography as a tool for antibody purification, Methods. 56 (2012) 116–129. https://doi.org/10.1016/j.ymeth.2011.10.007.

[2] M. Wilchek, T. Miron, Thirty years of affinity chromatography, React. Funct. Polym. 41 (1999) 263–268. https://doi.org/10.1016/S1381-5148(99)00042-5.

[3] J. Turková, O. Hubálková, M. KřivákováJ. Čoupek, Affinity chromatography on hydroxyalkyl methacrylate gels: I. Preparation of immobilized chymotrypsin and its use in the isolation of proteolytic inhibitors, Biochimica et Biophysica Acta (BBA) - Protein Structure. 322 (1973) 1–9. https://doi.org/10.1016/0005-2795(73)90167-0.

[4] E.L. Rodriguez, S. Poddar, S. Iftekhar, K. Suh, A.G. Woolfork, S. Ovbude, A. Pekarek, M. Walters, S. Lott, D.S. Hage, Affinity chromatography: A review of trends and developments over the past 50 years, J. Chromatogr. B Analyt. Technol. Biomed. Life Sci. 1157 (2020) 122332. https://doi.org/10.1016/j.jchromb.2020.122332.

[5] I.J. Bruce, J. Taylor, M. Todd, M.J. Davies, E. Borioni, C. Sangregorio, T. Sen Synthesis, characterisation and application of silica-magnetite nanocomposites, J. Magn. Magn. Mater. 284 (2004) 145–160. https://doi.org/10.1016/j.jmmm.2004.06.032.

[6] D. Tanyolaç, A.R. Özdural, A new low cost magnetic material: magnetic polyvinylbutyral microbeads, React. Funct. Polym. 43 (2000) 279–286. https://doi.org/10.1016/S1381-5148(99)00054-1.

[7] I. Safarik, M. Safarikova, Magnetic techniques for the isolation and purification of proteins and peptides, Biomagn. Res. Technol. 2 (2004) 7. https://doi.org/10.1186/1477-044X-2-7.

[8] I. Safarík, M. Safaríková, Use of magnetic techniques for the isolation of cells, J. Chromatogr. B Biomed. Sci. Appl. 722 (1999) 33–53. https://www.ncbi.nlm.nih.gov/pubmed/10068132.

[9] L. Chen, T. Wang, J. Tong, Application of derivatized magnetic materials to the separation and the preconcentration of pollutants in water samples, Trends Analyt. Chem. 30 (2011) 1095–1108. https://doi.org/10.1016/j.trac.2011.02.013.

[10] J.-H. Lin, Z.-H. Wu, W.-L. Tseng, Extraction of environmental pollutants using magnetic nanomaterials, Anal. Methods. 2 (2010) 1874–1879. https://doi.org/10.1039/C0AY00575D.

[11] J.H. Jung, J.H. Lee, S. Shinkai, Functionalized magnetic nanoparticles as chemosensors and adsorbents for toxic metal ions in environmental and biological fields, Chem. Soc. Rev. 40 (2011) 4464–4474. https://doi.org/10.1039/c1cs15051k.

[12] Y. Pan, X. Du, F. Zhao, B. Xu, Magnetic nanoparticles for the manipulation of proteins and cells, Chem. Soc. Rev. 41 (2012) 2912–2942. https://doi.org/10.1039/c2cs15315g.

[13] I. Sucholeiki, L.M. Toledo-Sherman, C.M. Hosfield, K. Boutilier, L.V. DeSouza, D.R. Stover, Novel magnetic supports for small molecule affinity capture of proteins for use in proteomics, Mol. Divers. 8 (2004) 9–19. https://doi.org/10.1023/b:modi.0000006780.36844.64.

[14] X.D. Tong, B. Xue, Y. Sun, A novel magnetic affinity support for protein adsorption and purification, Biotechnol. Prog. 17 (2001) 134–139. https://doi.org/10.1021/bp000134g.

[15] M. Shikida, K. Takayanagi, K. Inouchi, H. Honda, K. Sato, Using wettability and interfacial tension to handle droplets of magnetic beads in a micro-chemical-analysis system, Sens. Actuators B Chem. 113 (2006) 563–569. https://doi.org/10.1016/j.snb.2005.01.029.

[16] Y. Sasagawa, H. Danno, H. Takada, M. Ebisawa, K. Tanaka, T. Hayashi, A. Kurisaki, I. Nikaido, Quartz-Seq2: a high-throughput single-cell RNA-sequencing method that effectively uses limited sequence reads, Genome Biol. 19 (2018) 29. https://doi.org/10.1186/s13059-018-1407-3.

[17] T. Hayashi, H. Ozaki, Y. Sasagawa, M. Umeda, H. Danno, I. Nikaido, Single-cell full-length total RNA sequencing uncovers dynamics of recursive splicing and enhancer RNAs, Nat. Commun. 9 (2018) 619. https://doi.org/10.1038/s41467-018-02866-0.

[18] S. Picelli, Å.K. Björklund, O.R. Faridani, S. Sagasser, G. Winberg, R. Sandberg, Smart-seq2 for sensitive full-length transcriptome profiling in single cells, Nat. Methods. 10 (2013) 1096–1098. https://doi.org/10.1038/nmeth.2639.

[19] M. Hagemann-Jensen, C. Ziegenhain, P. Chen, D. Ramsköld, G.-J. Hendriks, A.J.M. Larsson, O.R. Faridani, R. Sandberg, Single-cell RNA counting at allele and isoform resolution using Smart-seq3, Nat. Biotechnol. 38 (6/2020) 708–714. https://doi.org/10.1038/s41587-020-0497-0.

[20] M. Hagemann-Jensen, C. Ziegenhain, R. Sandberg, Scalable single-cell RNA sequencing from full transcripts with Smart-seq3xpress, Nat. Biotechnol. 40 (2022) 1452–1457. https://doi.org/10.1038/s41587-022-01311-4.

[21] V. Hahaut, D. Pavlinic, W. Carbone, S. Schuierer, P. Balmer, M. Quinodoz, M. Renner, G. Roma, C.S. Cowan, S. Picelli, Fast and highly sensitive full-length single-cell RNA sequencing using FLASH-seq, Nat. Biotechnol. 40 (2022) 1447–1451. https://doi.org/10.1038/s41587-022-01312-3.

